# Hydrocarbon Metabolism and Petroleum Seepage as Ecological and Evolutionary Drivers for *Cycloclasticus*

**DOI:** 10.1101/2024.08.08.607087

**Authors:** Eleanor C. Arrington, Jonathan Tarn, Veronika Kivenson, Brook L. Nunn, Rachel M. Liu, Blair G. Paul, David L. Valentine

## Abstract

Aqueous-soluble hydrocarbons dissolve into the ocean’s interior and structure deep-sea microbial populations influenced by natural oil seeps and spills. *n-*Pentane is a seawater-soluble, volatile compound abundant in petroleum products and reservoirs and will partially partition to the deep-water column following release from the seafloor. In this study, we explore the ecology and niche partitioning of two free-living *Cycloclasticus* strains recovered from seawater incubations with *n*-pentane using and distinguish them as an open ocean variant and a seep-proximal variant, each with distinct capabilities for hydrocarbon catabolism. Comparative metagenomic analysis indicates the open ocean adapted variant encodes more general pathways for hydrocarbon consumption, including short-chain alkanes, aromatics, and long-chain alkanes, and also possesses redox versatility in the form of respiratory nitrate reduction and thiosulfate oxidation; in contrast, the seep variant specializes in short-chain alkanes and relies strictly on oxygen as the terminal electron acceptor. Both variants observed in our work were dominant ecotypes of *Cycloclasticus* observed during the Deepwater Horizon disaster, a conclusion supported by 16S rRNA analysis and read-recruitment of sequences from the submerged oil plume during active flow. A comparative genomic analysis of *Cycloclasticus* across various ecosystems suggests distinct strategies for hydrocarbon transformations among each clade. Our findings suggest *Cycloclasticus* is a versatile and opportunistic consumer of hydrocarbons and may have a greater role in the cycling of sulfur and nitrogen, thus contributing broad ecological impact to various ecosystems globally.

## Introduction

Much of the petroleum entering the ocean annually is introduced near the seafloor from human-caused incidents such as pipeline ruptures, well blowouts, and leaking submerged oil tankers, alongside other deep hydrocarbon inputs originating from natural oil seepage and hydrothermal vents. Following petroleum release to the seafloor, several compounds dissolve into seawater due to their aqueous solubility, subsequently affecting the microbial community within the surrounding water column [1–3]. These semi-aqueous soluble compounds can be overlooked as drivers of microbial metabolism in the deep community because these compounds often evaporate from surface oil slicks exposed to the atmosphere, which receive the majority of attention from agencies and scientists responding to oil-related incidents. This work focuses on the semi-aqueous-soluble compound *n-*pentane, which is known to partition to the deep ocean following release from the seafloor [2, 3].

Petroleum exposure to seawater drives a remarkable decrease in prokaryotic diversity due to a strong selection for hydrocarbon-degrading microorganisms and toxic effects on other taxa[4, 5]. Models of in-situ hydrocarbon biodegradation indicate that as a water parcel encounters a hydrocarbon source, a seed population of hydrocarbon degraders grows abundantly [6–11]. However, the origin and ecology of these seed populations are primarily hypothetical. Many studies have suggested seed populations are prolonged and sustained by hydrocarbon substrates originating from various sources, including hydrothermal vents [12], cyanobacteria, and eukaryotic phytoplankton[12–14], as well as natural gas seepage [15, 16]. As an example, the ubiquitous alkane degrader, *Alcanivorax*, exhibits basal cell populations that range from 10 to 5,000 cells per mL in uncontaminated seawater [6, 17], with recent work suggesting high native abundance is subsidized by widespread biosynthesis of long-chain *n*-alkanes by marine phytoplankton [13, 18]. Recent evidence has shown that methanotrophs can be physically transported on bubbles from a gas seep [19, 20], pointing to seeps as a physical mechanism to seed the water column with hydrocarbon degraders. Alternatively, facultative hydrocarbon degraders could be present that rely on other metabolic inputs such as amino acids, carbohydrates, lipids, or other organic acids and switch to hydrocarbons under appropriate conditions [21–23]. Very few studies have focused on how these factors control the development of a petroleum-degrading community during oil spills in previously uncontaminated waters.

Our investigation [18]of the ocean’s biological hydrocarbon cycle revealed the microbial response to *n-*pentane is structured by proximity to seepage (Fig. 1). This previous work and our current study focus on sea-going incubations conducted with water collected from the deep ocean (1 000 m) along a transect spanning the Gulf of Mexico (GOM) and the North Atlantic. *n-*Pentane metabolism was observed through a closed-system optical oxygen technique in which population blooms were defined by three consecutive time points of oxygen loss greater than 0.21 µM h^−1^. We observed distinct bloom response times to *n*-pentane in relation to natural seepage, whereby bloom onset on pentane is ∼9X faster in the seep-ridden Northwest Gulf of Mexico compared to the water underlying the North Atlantic subtropical gyre [18]. Median bloom times varied from 72.9 days furthest from natural seepage to 8.3 days closest to natural seepage. The fraction of samples that exhibited a bloom response also coincided with proximity to natural seepage (Fig. 1b).

**Figure 1.**
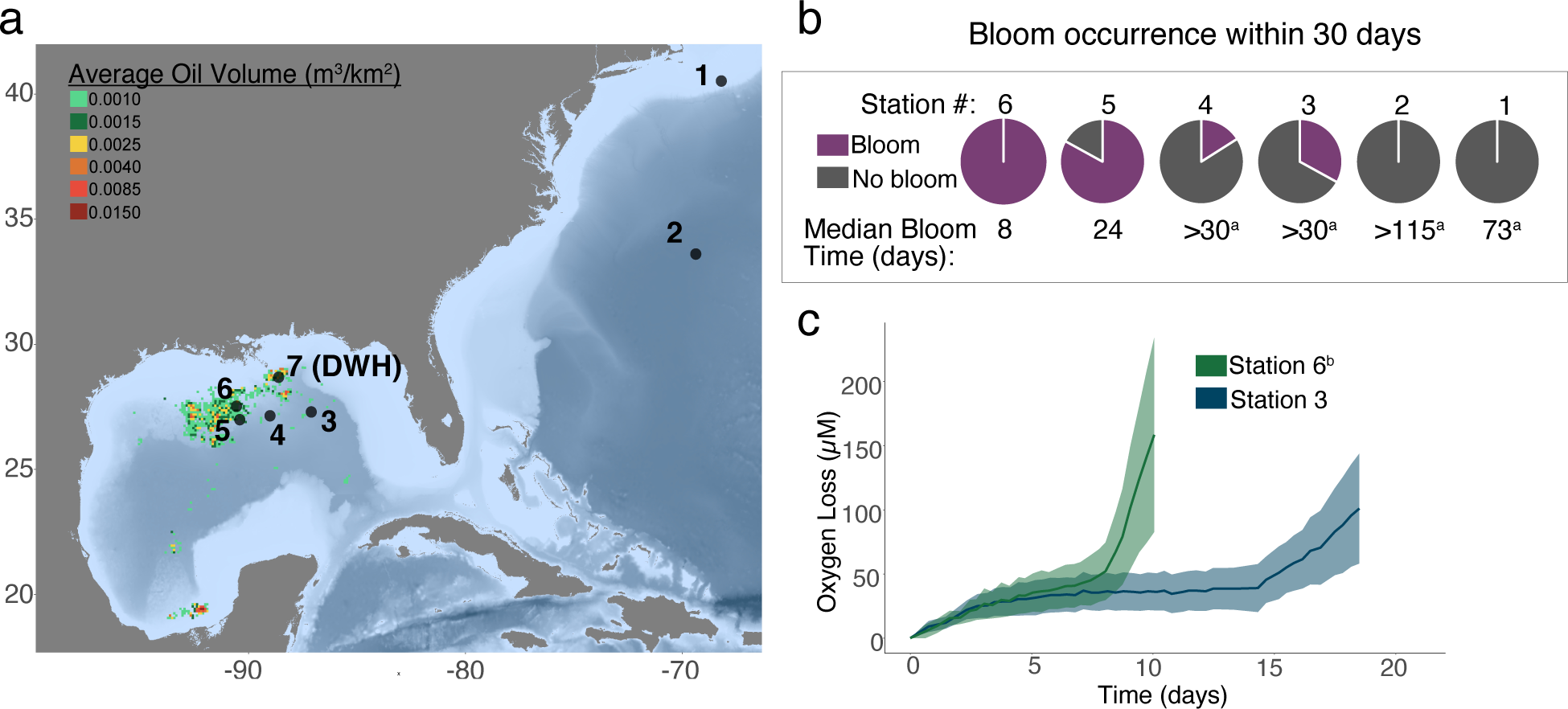
Previously published data in [18]. Pentane biodegradation informed by respiration after normalization to unamended controls. **a** Sampling stations relative to natural petroleum seepage. Seep locations from [60]. DWH indicates the location of the sample collected during the DWH disaster. **b** Fraction of incubations that bloomed and bloom timing. Data previously published [18]. **c** Oxygen loss in pentane incubations with time for blooms at station 6 vs station 3. ^b^ indicates that one bloom (sample 180) was omitted as it was an outlier with delayed bloom response (23 days) compared to the other five replicates which bloomed in 8 days.

Here, we investigate the influence of biogeography on microbial hydrocarbon metabolism by analyzing the genomic content of organisms with contrasting bloom response and source water origins. We extract and analyze blooms’ 16S rRNA amplicon content, assemble high-quality draft genomes from bloom experiments, and perform a complementary proteomic analysis to evaluate metabolism. The predominant microbial member of the *n-*pentane-enriched community across the Gulf of Mexico belongs to the *Cycloclasticus* genus, originally named for their metabolic capability to consume polycyclic aromatic hydrocarbons (PAHs). Interestingly, we observe two strains of *Cycloclasticus* bloom in response to short-chain alkanes in the Gulf of Mexico, with one strain favoring the seep-influenced region of the northwestern GOM (hereafter referred to as the seep variant, SV). In contrast, the other strain favors the open ocean region far from natural seepage (hereafter referred to as the open ocean variant, OOV).

## Methods

### Incubation design and sample collection

Seawater samples were collected on two research cruises aboard RV Atlantis in June 2015 and the RV Neil Armstrong in May 2017. *n-*Pentane incubations were conducted at stations 1 (40° 9.14ʹ N, 68° 19.889ʹ W), 2 (33° 58.21ʹ N, 69° 43.38ʹ W), 3 (27° 30.41ʹ N, 87° 12.41ʹ W), 4 (27° 15.00ʹ N, 89° 05.05ʹ W), 5 (27° 11.60ʹ N, 90° 41.75ʹ W) and 6 (27° 38.40ʹ N, 90° 54.98ʹ W) with seawater collected from 1 000 m. Respiration data and methods are available from [18], with sample numbers re-named for this study to exclude irrelevant data. Seawater collected from the CTD Niskin bottles was transferred to 250 mL glass serum vials using a small length of Tygon tubing. Vials were filled for at least 3 volumes of water to overflow. Care was taken to ensure no bubbles were present before sealing with a polytetrafluoroethylene (PTFE) coated chlorobutyl rubber stopper and crimp cap seal. All bottles, except for unamended blank controls, immediately received 10 μL of pentane using a gas-tight syringe (Hamilton, Nevada, USA) and were maintained in the dark at in-situ temperature (4 °C). Before filling, each serum bottle was fixed with a contactless optical oxygen sensor (Pyroscience, Aachen, Germany) on the inner side with silicone glue, and afterward were cleaned from organic contaminants with triple rinses of ethanol, 3% hydrogen peroxide, 10% hydrochloric acid, and MilliQ water, and were sterilized via autoclave. Oxygen concentration was monitored approximately every 8 hours with a fiber optic oxygen meter (Pyroscience, Aachen, Germany). Observed changes in oxygen content were normalized to unamended controls to correct for oxygen loss from background respiration processes and variability due to temperature changes.

### DNA extraction, 16S rRNA community analysis

DNA was only analyzed from stations within the Gulf of Mexico. After 27-30 days, each incubation was harvested and filtered on a 0.22 µm polyethersulfone filter. DNA extraction was performed from ¼ of each filter using the PowerSoil DNA extraction kit with the following modifications: 200 µl of bead beating solution was removed at the initial step and replaced with phenol-chloroform, the C4 bead binding solution was supplemented with 600 µL of 100% ethanol, and we added an additional column washing step with 650 µL of 100% ethanol. Extracts were purified and concentrated by ethanol precipitation, then stored at -80 °C. The V4 region of the 16S rRNA gene was amplified using the method described by [24] with small modifications to the 16Sf and 16Sr primers according to[25, 26]. Amplicon PCR reactions contained 1 µL of template DNA, 2 µL of forward primer, 2 µL of reverse primer, and 17 µL of AccuPrime Pfx SuperMix. Thermocycling conditions consisted of 95° 2 min, 30 cycles of 95°C for 20 secs, 55°C for 15 secs, 72°C for 5 min, and a final elongation at 72°C for 10 min. Sample DNA concentrations were normalized using the SequelPrep Normalization Kit, cleaned using the DNA Clean and Concentrator kit, visualized on an Agilent Tapestation, and quantified using a Qubit Flurometer. Samples were sequenced at the UC Davis Genome Center on the Illumina MiSeq platform with 250nt, paired-end reads. A PCR-grade water sample was included in extraction, amplification, and sequencing as negative control to assess for DNA contamination.

Trimmed fastq files were quality filtered using the fastqPairedFilter command within the dada2 R package, version 1.9.3 (Callahan et al., 2016) with following parameters: truncLen=c(190,190), maxN=0, maxEE=c(2,2), truncQ=2, rm.phix=TRUE, compress=TRUE, multithread=TRUE. Quality filtered reads were dereplicated using derepFastq command. Paired dereplicated fastq files were joined using the mergePairs function with the default parameters. A single nucleotide variant (SNV) table was constructed with the makeSequenceTable command and potential chimeras were removed denovo using removeBimeraDenovo. Taxonomic assignment of the sequences was done with the assignTaxonomy command using the Silva taxonomic training dataset formatted for DADA2 v132 [27, 28]. If sequences were not assigned, they were left as NA.

### Metagenomic sequencing and reconstruction

Metagenomic library preparation and shotgun sequencing were conducted at the University of California Davis DNA Technologies Core. DNA was sequenced on the Illumina HiSeq4000 platform, producing 150-base pair (bp) paired-end reads with a targeted insert size of 400 bp. Quality control and adaptor removal were performed with Trimmomatic [29] (v.0.36; parameters: leading 10, trailing 10, sliding window of 4, quality score of 25, minimum length 151 bp) and Sickle [30] (v.1.33 with paired-end and Sanger parameters). 10-30% of the trimmed high-quality reads were randomly subsampled to deconvolute assembly and downstream binning in samples with very high coverage. The subsampled high-quality reads were assembled using metaSPAdes [31] (v.3.8.1; parameters *k* = 21, 33, 55, 77, 88, 127). The quality of assemblies was determined using QUAST [32] (v.5.0.2 with default parameters). Sequencing coverage was determined for each assembled scaffold by mapping high-quality reads to the assembly using Bowtie2 [33] (v.2.3.4.1; default parameters) with Samtools[34] (v.1.7). Contigs greater than 2,500 bp were manually binned using Anvi’o with Centrifuge (v.1.0.1) based on coverage uniformity (v.5) [35, 36]. Quality metrics for metagenome-assembled genomes (MAGs) were determined using CheckM [37] (v.1.0.7; default parameters). The taxonomy of each MAG was classified using GTDB-Tk (v.1.0.2) against The Genome Taxonomy Database [38] (https://data.ace.uq.edu.au/public/gtdb/data/releases/release89/89.0/, v.r89). The average nucleotide identity of each genome was determined with the ANI Matrix via the Enveomics tool collection[39]. For the DWH metagenomes, the second variant related to OOV could not be recovered with metagenomics due to issues with assembly fragmentation and binning of the two closely related strains. To obtain a high-quality draft genome of “MAG_DWH_1”, we subsampled our reads by 50% until the OOV-related sequences were a small fraction of the assembled data.

Using a 16S search tool through the Joint Genome Institute-Integrated Microbial Genomes and Microbiomes (JGI-IMG) portal, we identified public unprocessed environmental metagenomic datasets with *Cycloclasticus* representation. These datasets were downloaded, and metagenomic reconstruction was performed according to the above protocol with the following modifications: binning was performed using the automated binning software MetaBAT2 [40]. We have received explicit permission from the authors of each dataset to include their data in our analyses.

### Phylogenetics

To define genome phylogenomic relationships of MAGs, 16 universal ribosomal proteins (RPs) were used L2-L6, L14-L16, L18, L22, L24, S3, S8, S10, S17, and S19. For phylogenies of metabolic genes and ribosomal proteins, all representative sequences and concatenated alignments containing <25% informative sites were excluded in tree construction. Each protein was aligned using MUSCLE (v.3.8.425) [41]. All columns with >95% gaps were removed using TrimAL[42]. Maximum-likelihood phylogenetic analysis of concatenated alignment was inferred by RAxML[43] (v.8.9; parameters: raxmlHPC -T 4 -s input -N autoMRE -n result -f a -p 12345 -x 12345 -m PROTCATLG). Resulting trees were visualized using FigTree [44] (v.1.4.3).

### Metaproteomics

We analyzed metaproteomes from two of the SV (seep variant) samples with corresponding metagenomes (“3_C5_1” and “3_C5_2”) as the reference databases. Proteins from each sample were extracted and prepared from ¼ filter (equivalent to ∼60 mL of filtered water and ∼1.3×10^8^ bacteria) for liquid chromatography and tandem mass spectrometry (LC-MS/MS) using a protocol adapted from [45]. Briefly, filters were cut into 2 mm pieces and submerged in 100 µL of 6M urea and 600 µL of 50 mM NH4HCO3 and sonicated (Branson 250 Sonifier; 20 kHz, 5 × 20 s on ice). to lyse cells. Protein concentrations for each sample were quantified in triplicate using a Bicinchoninic Acid protein assay kit (Pierce Thermo Scientific) using a microplate reader. Proteins within the lysate were reduced and alkylated using dithiothreitol and iodoacetamide, respectively, digested with Trypsin (12 h; 1:20 enzyme to protein) and desalted with C18 centrifugal spin columns. Peptides were dried down and resuspended in 2% ACN, 0.1% formic acid before analysis with a nanoAcquity UPLC (Waters Corp., Milford, MA) in line with a Q-Exactive-HF (Thermo Fisher Scientific, Waltham, MA). Reverse phase chromatography was achieved using a PicoTip (New Objective) fused silica capillary column (75 μm i.d., 30 cm long) packed with C18 beads (Dr. Maisch ReproSil-Pur; C18-Aq, 120 Å, 3 μm). The analytical column was preceded by a 150 μm i.d. PicoFrit (New Objective) precolumn packed with C-18 beads to 3 cm long (Dr. Maisch ReproSil-Pur; C18-Aq, 120 Å, 3 μm). Peptides were eluted using a 90-minute acidified (formic acid, 0.1% v/v) water-acetonitrile gradient (2-45% acetonitrile). Sample analyses on the MS were randomized to reduce batch effects, and quality control (QC) peptide mixtures were analyzed every six injections to monitor chromatography and MS sensitivity. Each sample was analyzed with data-dependent acquisition (DDA). From precursor ion scans of 400–1200 m/z the top 20 most intense ions were selected for MS2 acquisition. Centroid full MS resolution data was collected at 70,000 with AGC target of 1 ×106 and centroid MS2 data was collected at resolution of 35,000 with AGC target of 5 × 104. Dynamic exclusion was set to 20 seconds and +2, +3, +4 ions were selected for MS2 using DDA mode. Comet [46, 47] was used to search the DDA files against the 2 metagenomes concatenated with 50 common contaminants and the QC peptides. Comet parameters included: 10 ppm precursor mass tolerance, fully-tryptic specificity with 0 allowed missed cleavages, cysteine modification of 57 Da and modifications on methionine of 15.999 Da. PeptideProphet was used to validate peptide spectral matches (PSMs) and determine thresholds for a false discovery rate [48]. All relevant peptide hits with an e-value less than 0.01 were used to define protein presence. All peptides’ tandem mass spectra discussed here were manually investigated to verify b and y ions. The mass spectrometry data are deposited to the Proteome Xchange Consortium via the PRIDE partner repository with the dataset identifier PXD022428.

### Genomic Read Mapping

To evaluate the prevalence of *Cycloclasticus* MAGs and single amplified genomes (SAGs) in the newly sequenced sample from the Deepwater Horizon event, we used CoverM (v.0.7.0) [49] the bwa-mem read-mapping parameter with a minimum identity of >95% over >95% of the read length.

## Results and Discussion

### Variant biogeography

Incubations conducted with seawater from 1,000 m depth containing ambient nutrients successfully exhibited robust blooms of bacteria when supplied with *n-*pentane as a carbon and energy source. Blooms are characterized by bacterioplankton abundance, increasing by ∼10X (Supplementary Dataset S1), significant drawdowns in inorganic phosphate and nitrate concentration (Fig. 2c, Fig. 2d, Fig. 2e; Supplementary Dataset S1), and the emergence of dominant taxa comprising >70% of the microbial community at the termination of the experiment (Fig 2a). Community analysis of incubations that failed to bloom within the experimental timeframe (27-30 days) revealed instances where the community was dominated by a limited number of taxa, indicating a microbial community shift precedes major respiratory signals (Supplementary Dataset S2, Supplementary Information Fig. S3). Over 60% of all pentane blooms in the deep GOM were dominated by the *Cycloclasticus* genus (Fig 2a). Variation among replicate incubations was observed as a change in the dominant taxa, which occurred more often at stations closer to natural seepage, potentially related to greater microbial diversity and abundance of alkane degraders in those waters. At station 6, *Colwellia* and an unclassified genus belonging to *Cellvibrionaceae* bloomed, and the same unclassified *Cellvibrionaceae* bloomed at station 5 (Fig 2a).

**Figure 2.**
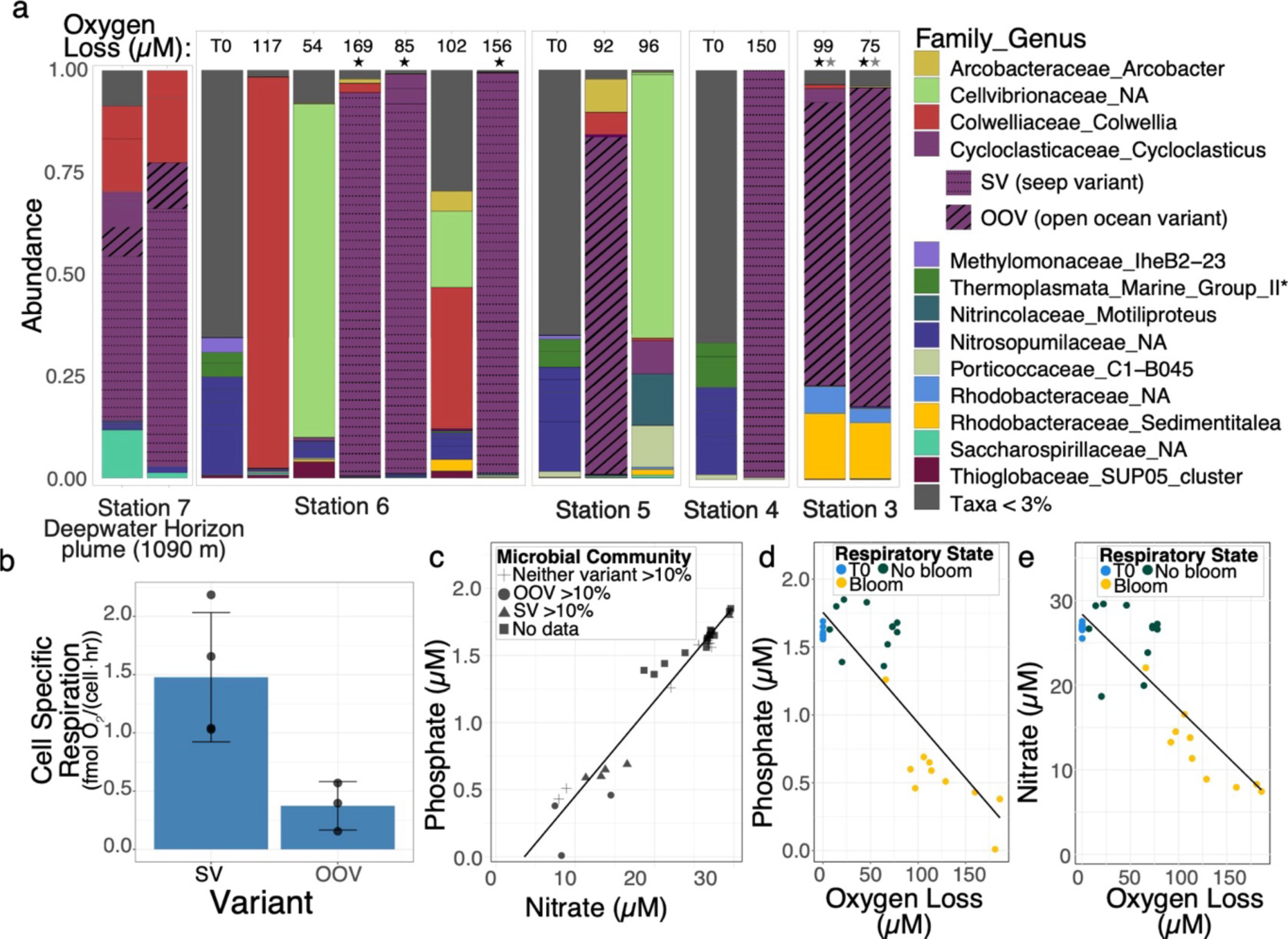
Microbial and geochemical characteristics of blooms. **a** Microbial community composition of pentane blooms informed via the V4 region of the 16S rRNA gene for initial environmental samples (denoted “T0”) and pentane incubations harvested after significant oxygen loss was observed. DWH sample collected on 5/30/10 during the DWH event. SV strain *Cycloclasticus* blooms in 8-20 days in locations close to natural seepage and was abundant on 5/30/10 during the Deepwater Horizon disaster. OOV strain of *Cycloclasticus* blooms in 16-20 days and dominates seawater originating further from natural seepage inputs. *No taxonomic representative at the family or genera level. “Oxygen loss” indicates the change in oxygen relative to initial concentrations when DNA sample was collected **b** Cell specific respiration in incubations dominated (>80%) by SV and OOV *Cycloclasticus*. **c** Dissolved phosphate vs dissolved nitrate concentration in initial samples (T0) and at sample harvest. **d** Dissolved phosphate concentration in final samples is depleted compared to unamended controls and initial samples (T0). **e** Dissolved nitrate concentration in final samples is depleted compared to unamended controls and initial samples (T0).

Among blooms, we observed two dominant *Cycloclasticus* variants, with one (SV) being the primary blooming population at station 6 within the northwestern GOM seep field and the other (OOV) being the primary blooming organism at station 3 within the more petroleum-depleted region (Fig. 1a, Fig. 2a, Supplementary Data S3). The occurrence of SV and OOV at stations 4 and 5 is patchier, with both variants occurring at station 4 and only OOV blooming at station 5 (Fig. 2a). The success of OOV over SV at station 5 may be explained by the western flow of eastern sourced seep-deplete waters observed at the time seawater was collected (Supplementary Data Fig. S7). Cell-specific respiration is higher for SV than the OOV (Fig. 2b), and the respiration profile (Fig. 1c) of these variants also showed distinct patterns, where OOV presents more gradual oxygen consumption with time. The OOV was also observed in small relative abundances (<1%) at station 6 (Supplementary Dataset S2). In each *n-*pentane incubation where the two variants co-occurred, the SV numerically dominated by ∼3 orders of magnitude except for the *Colwellia*/*Cellvibrionaceae* bloom at station 6, where both variants were detected at <1% abundance. This suggests that SV is better adapted to conditions associated with natural seepage, as it outcompeted OOV, which only blpoomed at stations that are distant from seeps.

### Pentane metabolism

We reconstructed high-quality metagenomes from five pentane bloom samples, with completeness >97% and redundancy <2% (black stars in Fig 2). Three MAGs, named “6_C5_1”, “6_C5_2”, and “6_C5_3”, originated from station 6 (natural seep region), and two MAGs, named “3_C5_1” and “3_C5_2”, originated from station 3 (open ocean region). Our analyses were further supported by proteomic analysis of the two MAGs from station 3 (gray stars in Fig. 2a; Fig 4b). Each MAG was recovered from biologically independent incubations, yet every significant component of metabolism and taxonomic marker analyzed was nearly identical within each ecotype variant; therefore, we will refer to “SV-MAG” as the three MAGs from station 6 and “OOV-MAG” as the two MAGs from station 3. Within both SV- and OOV-MAGs, we found genomic potential for *n-*pentane utilization for catabolism and anabolism (Fig. 3).

**Figure 3.**
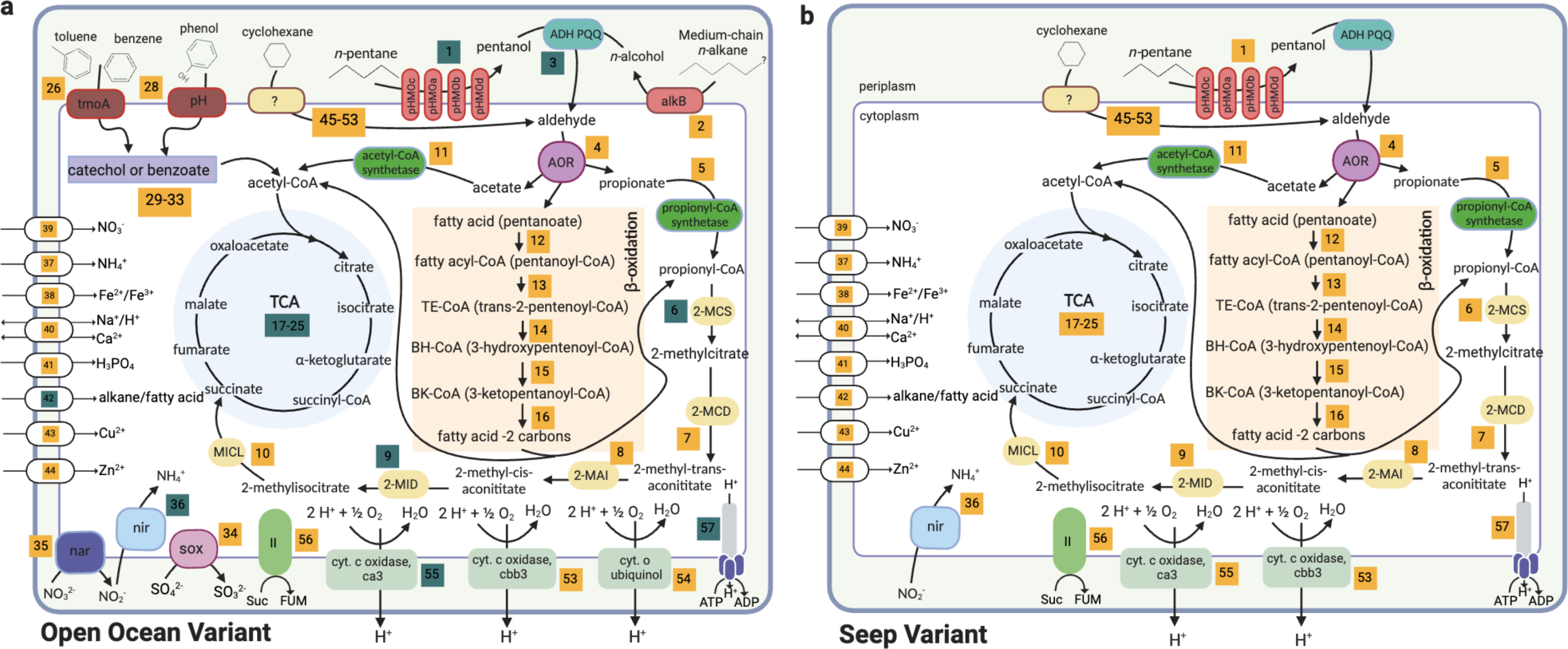
Carbon, nitrogen, and sulfur metabolism present in *Cycloclasticus* variants blooming on pentane. **a** Left cell indicates OOV-MAG, and **b** depicts SV-MAG. Yellow boxes indicate a reaction (and its reference number) that could be linked with a predicted metabolic function, see Supplementary Dataset S3. Blue boxes indicate peptides for that enzyme were observed in proteomic data. Proteomics performed on only OOV-MAG. If the reaction box describes multiple enzymes, only one needs to be observed in proteomic data for it to be colored blue. Enzyme abbreviations: particulate hydrocarbon monooxygenase pHMO (a,b,c,d); PQQ-dependent alcohol dehydrogenase (ADH PQQ); aldehyde oxidoreductase (AOR); 2-methylcitrate synthase (2-MCS); 2-methylcitrate dehydratase (2-MCD); 2-methylcitrate isomerase (2-MAI); 2-methylisocitrate dehydratase (2-MID); methylisocitrate lyase (MICL); nitrite reductase (Nir); respiratory nitrate reductase (Nar); thiosulfate oxidation complex (Sox); alkane-1-monooxygenase (alkB); toluene monooxygenase (tmoA); phenol/toluene 2-monooxygenase (pH). The tricarboxylic acid (TCA) and beta-oxidation pathway are highlighted in blue and peach colors.

**Figure 4.**
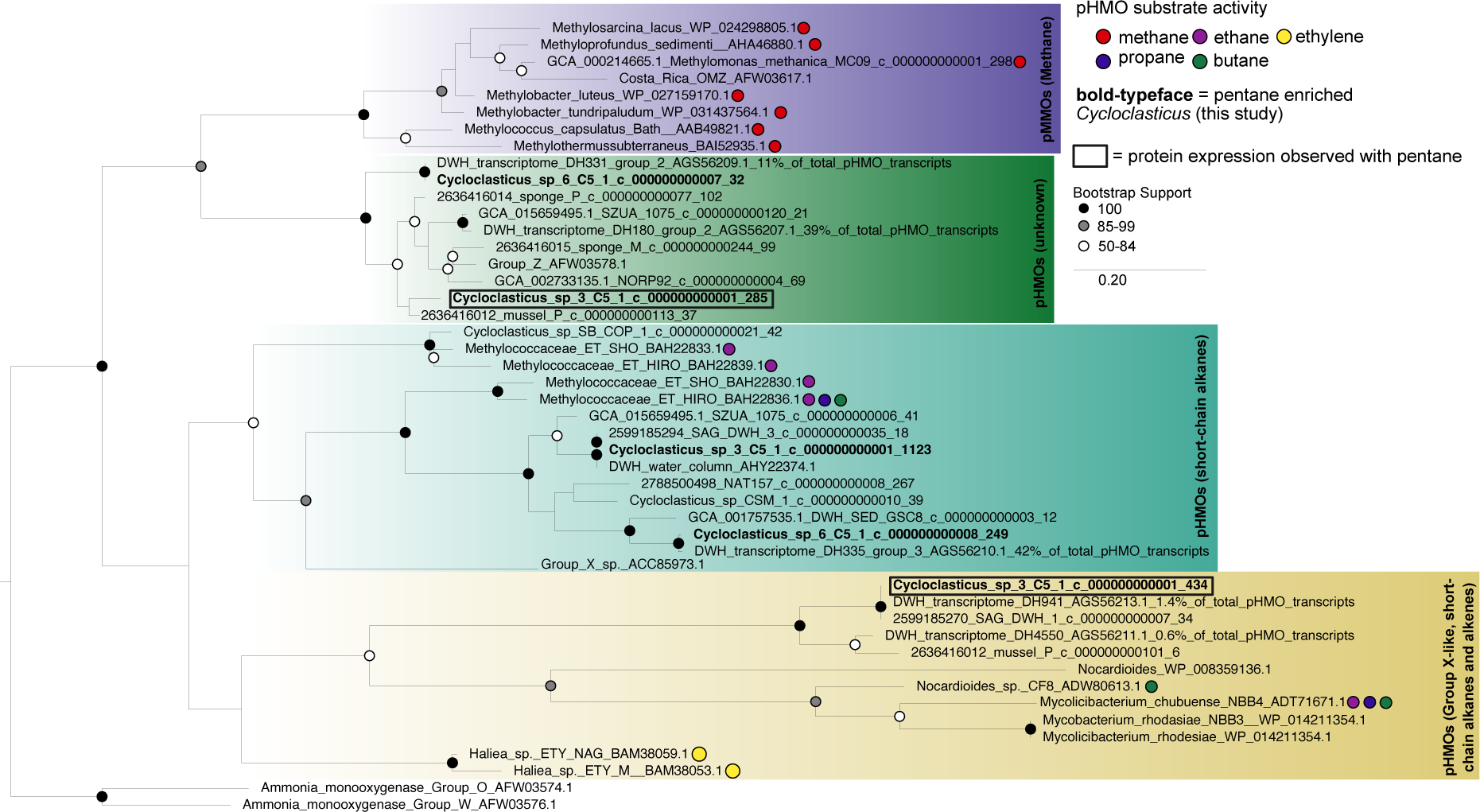
Maximum likelihood tree of pHMO subunit A drawn to scale, with branch lengths representing the number of substitutions per site. Bootstrap values below 50% are not shown. Each major clade is color coded for readability with purple representing pMMOs with activity on methane, the green clade is “unknown” and lacks any known substrate specificity, the cyan clade represents group X pHMOs (ethane/ethylene, propane, and butane activity), and the yellow clade are group-X like (ethane, propane, and butane activity). Sequences from the pentane enriched *Cycloclasticus* MAGs are in bold, boxed values indicate pHMOs detected in proteomic data.

The first step in the consumption of pentane is the oxidation to pentanol, and we hypothesize that this step is catalyzed by the copper-containing membrane-associated monooxygenase, called particulate hydrocarbon monooxygenases (pHMOs). The most well-characterized pHMO is the particulate methane monooxygenase, which oxidizes methane to methanol [50]. We found multiple copies of genes encoding pHMOs in both *Cycloclasticus* MAG variants (Fig. 3, Fig. 4, Supplementary Information Fig. S8). Each copy of pHMO varies phylogenetically from the other copies within the same genome, suggesting each operon may have different substrate specificities or capitalize on alkanes of varying substrate concentrations [51] (Fig. 4). Both MAG-SV and MAG-OOV have pHMO sequences that form monophyletic clades with reference sequences with demonstrated affinity for ethane and butane. Both variants also contain a sequence that forms a monophyletic clade that is only distantly related to pMMOs; however, this clade contains no currently validated reference sequences, and we refer to its function as “unknown”. Proteomics confirmed the expression of pHMOs, specifically subunit a and b (Fig. 3, Fig. 4) in the presence of pentane. The two pHMOs for which peptides were detected in OOV belongs to a sequence from OOV-MAG in the “unknown” clade of pHMOs and one clade containing reference sequences with demonstrated affinity for ethane and butane. *AlkB*, a gene known to function on medium to long-chain alkanes, was found encoded in the MAG-OOV; however, no peptides were observed in the proteomics analyses (Fig. 3). Still, given the minimal sample size analyzed for proteomics and the potential for false negatives due to e.g. ionization and extraction efficiencies, we do not exclude the possibility that *alkB* could also be active in these samples and used to consume pentane by the OOV.

The second step in the consumption of pentane is the conversion of pentanol to an aldehyde. In many bacteria that oxidize alcohols, this reaction is catalyzed by pyrroloquinoline quinone-dependent alcohol dehydrogenases (PQQ-ADH). We found genes encoding PQQ-ADHs in both *Cycloclasticus* MAG variants and proteomic expression of PQQ-ADH in OOV-MAG samples. (Fig. 3, Supplementary Information Fig S2). None of the PQQ-ADHs formed a monophyletic clade with reference sequences known to act on methanol, providing evidence against methane metabolism in SV and OOV. In the third step of pentane consumption, the aldehydes are oxidized to carboxylic acids, which could be achieved via a tungsten-containing aldehyde ferredoxin oxidoreductase (AOR), known to use short-chain alkane-derived aldehydes as their substrate [15, 52]. This conversion can also be performed by PQQ-ADH, as activity on aldehydes has been confirmed with reference sequences related to those encoded by SV and OOV (Supplementary Information Fig S2). Here, pentanoate is likely beta-oxidized using acyl-CoA dehydrogenase and enoyl-CoA hydratase and shunted into central carbon metabolism via the citric acid cycle (Fig. 3).

### Differences in variant metabolic potential

Interestingly, the metabolic capabilities of the SV-MAG and OOV-MAG differ substantially (Fig. 3, Fig. 4). The OOV-MAG encodes for general hydrocarbon metabolism that includes the nearly complete pathway for toluene consumption via the toluene monooxygenase conversion of toluene to benzoate (7 of 8 genes), benzoate conversion to catechol (3 of 4 genes), and the catechol metacleavage to acetyl-CoA which enters the tricarboxylic acid cycle (13 of 13 genes). The OOV-MAG also encodes toluene 2-monooxygenase, which converts benzene to catechol (6 of 6 genes) that can also be shunted through the same catechol meta-cleavage pathway as toluene to form acetyl-CoA (13 of 13 genes) and enter the tricarboxylic acid cycle. The OOV could also use the toluene-2 monooxygenase system to convert toluene to 3-methylcatechol (6 of 6 genes) and then convert 3-methylcatechol to acetyl-CoA and shunt to the tricarboxylic acid cycle (3 of 5 genes). Furthermore, the OOV-MAG encodes for *alkB* (1 of 1 gene), which is commonly used by other organisms for consumption of long-chain alkanes via beta-oxidation (OOV encodes 7 of 7 genes), resulting in propionyl-CoA and acetyl-CoA, which are also incorporated into the tricarboxylic acid cycle. Interestingly, neither OOV or SV (or any other *Cycloclasticus* genome analyzed in this study) encode a complete canonical naphthalene degradation pathway (naphthalene 1,2, dioxygenase is missing from all genomes), yet the strain *Cycloclasticus* SP-1 has been experimentally validated to use naphthalene as a sole carbon source (Fig. 5b) [53]. *Cycloclasticus* SP-1 and OOV-MAG encode 3 of 10 genes for naphthalene degradation, which indicates that OOV-MAG can also likely metabolize naphthalene, whereas SV encodes 0 of 10 genes.

**Figure 5.**
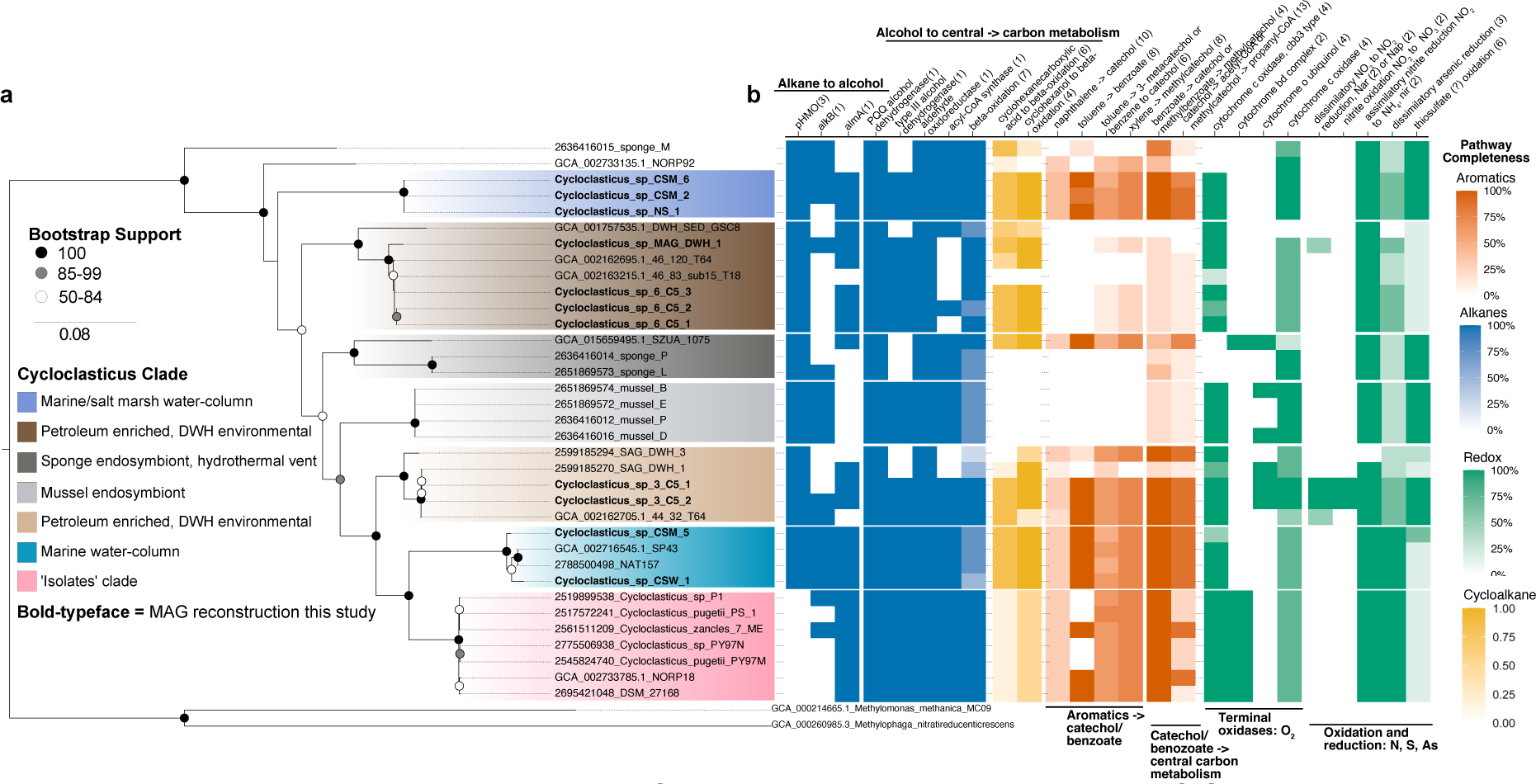
Phylogeny and capabilities for hydrocarbon consumption of *Cycloclasticus*. **a** Phylogeny constructed using 16 ribosomal proteins, subclades are designated based on relative distance from the root and supported by %ANI (Supplementary Information, Figure S5). **b** Completeness of select metabolic pathways relating to alkane (blue), aromatic (red), cycloalkane (yellow) metabolism and, redox (green). The number of genes considered in calculating pathway completeness are shown in parentheses.

Overall, the SV-MAG lacks many metabolic pathways for longer-chain alkanes and aromatic compounds compared to the OOV-MAG, seemingly limiting its hydrocarbon metabolism potential (Fig. 4, Fig. 5b). These observed differences are consistent with SV specialization on short-chain, aqueous-soluble alkanes and biogeography that includes seeding from the petroleum-rich source region in the Northern Gulf of Mexico. The genomic capacity for catabolism of multiple hydrocarbon classes in the OOV-MAG is consistent with an ability to capitalize on diffuse hydrocarbon sources that expand beyond natural seeps, including atmospheric deposition, terrestrial runoff, biogenic inputs, and oil spills. This enhanced capacity in OOV is consistent with an expanded biogeographic range relative to SV, which appears to be more highly reliant on substrate sourced from natural seepage.

### Anaerobic metabolism in *Cycloclasticus*

Anaerobic metabolism has yet to be observed in *Cycloclasticus,* and it remains unknown how these bacteria could contribute to hydrocarbon cycling in oxygen minimum zones or anoxic sediments. Here, we show the OOV variant of *Cycloclasticus* exhibits adaptations for life without oxygen, including the occurrence of genes for respiratory nitrate reductase (*Nar*), as well as a potential linkage to thiosulfate metabolism (Fig. 4). In OOV, we identify a complete canonical *nar* operon (*narGHJI*) encoding: i) the α subunit that catalyze NO3^−^ reduction to NO2^−^ (*narG*); ii) the iron–sulfur-containing β subunit (*narH*) that transfers electrons to the molybdenum cofactor of *narG*; iii) the *narJ* chaperone involved in enzyme formation and iv) the transmembrane cytochrome *b*-like γ subunit (*narI*) involved in electron transfer from membrane quinols to *narH*. Phylogenetic placement of *Cycloclasticus narG s*equences also confirms the relation to *narG* reference sequences (Supplementary Information Fig. S1). OOV also contains the *sox* operon (*soxCDXYZAB*), which encodes periplasmic sulfur-oxidizing proteins (Fig. 4). This operon can be used as a means of detoxification in some Gammaproteobacteria [8]; however, we do not exclude the possibility that *Cycloclasticus* could employ a lithoheterotrophic strategy. The use of thiosulfate to supplement heterotrophy is a strategy that has been demonstrated in other proteobacteria and could be useful in seeps and other benthic environments [54]. It is unclear how members of *Cycloclasticus* may access *n-*pentane in the absence of oxygen. No enzymes related to alkyl succinate synthase were detected. Multiple putative hits for the DMSO protein superfamily were detected, and this superfamily encompasses a variety of functions, including the anaerobic alkane degradation enzyme, alkane C2 methylene hydroxylase; however, OOV-MAG sequences do not form a monophyletic clade with reference sequences of this function (data not shown).

### Deepwater Horizon *Cycloclasticus*

The microbial response to the 2010 Deepwater Horizon blowout in the Gulf of Mexico induced blooms of *Cycloclasticus* in the deep ocean from large-scale intrusions of dissolved hydrocarbons [55]. These DWH blooms included multiple *Cycloclasticus* 16S rRNA sequence variants [56], which led us to ask whether SV and OOV were among those DWH variants. Therefore, we extracted and analyzed two replicate archive DNA samples collected while the wellhead was still leaking into the Gulf of Mexico on 5/30/10 from a depth of 1,090 m. At this depth, there was an oxygen anomaly characteristic of the respiratory response associated with the DWH subsurface intrusions [57] (Supplementary Information Fig. S6). Upon initial analysis of the microbial community via the V4 region (252 bp) of the 16S rRNA gene, we found the SV-MAG to be identical to the dominant member of the DWH sample and OOV-MAG to be identical to the second most abundant *Cycloclasticus* 16S rRNA single nucleotide variant (Fig. 2a).

We carried out additional shotgun sequencing on DNA from one of the DWH samples analyzed for 16S rRNA community analysis that was collected during the event. [15]Using read-mapping, we find that the fraction of the SV-MAG covered by the DWH reads spans 98% of the genome with approximately 20X coverage. The OOV-MAG is 95% covered from reads mapped from the DWH sample with approximately 5X coverage. We reconstructed a high-quality metagenome, here named “MAG_DWH_1”, which is 94% complete and 3.3% redundant. Upon expanding our analysis to the full-length 16S rRNA gene (as opposed to the V4 region in Fig 2), we find that the SV-MAG is 99.5% identical to MAG_DWH_1. Through a phylogenomic analysis of 16 ribosomal proteins, we find MAG_DWH_1 forms a monophyletic clade with SV-MAG (Fig. 5a, Supplementary Information Fig. S4). For comparison, we also drew from our previously published single amplified genomes (SAGs) from DWH, which are 71%, 49%, and 46% complete and herein referred to as “SAG_DWH_3”, “SAG_DWH_1”, and “SAG_DWH_2” [15]. We find that “SAG_DWH_1” and “SAG_DWH_3” are closely related to OOV-MAG, whereas “SAG_DWH_2” appears to be related to SV-MAG (Supplementary Information Fig. S4). For the relation of SV and OOV to the SAGs and MAG_DWH_1, we also find supporting evidence in the analysis of Average Nucleotide Identity and the 16S rRNA phylogeny (Supplementary Information Fig. S4 and S5). These results indicate a previously unrecognized distinction in the microbial response to the DWH event – that SV-like *Cycloclasticus* may have responded specifically to the highly abundant soluble *n*-alkanes. In contrast, OOV-like *Cycloclasticus* may have responded to soluble *n*-alkanes and other components, including benzene and toluene.

To further assess the ecological relevance of SV and OOV *Cycloclasticus* to DWH, we compared the similarities in the pHMO phylogenetic placement between SV- and OOV-MAGs and previously published transcripts from DWH subsurface plumes (Fig. 4) [58]. These results indicate that pHMOs most closely related to SV and OOV *Cycloclasticus* were expressed at high relative abundance during the DWH event, consistent with data showing the rapid microbial response by *Cycloclasticus* to short-chain n-alkanes (but not methane) concurrent with active discharge[55]. The pulse of bacterial growth in the deep ocean from the DWH event has been estimated at > 10^23^ cells, with a substantial fraction being SV *Cycloclasticus*. We, therefore, questioned if this level of ecological disturbance might have structured the hydrocarbon-degrading community in the GOM through 2015 when samples for this work were collected. More data is needed to assess this hypothesis rigorously. However, we note that methanotrophic biomass remained elevated in the years following the DWH event, perpetuating elevated methanotrophic activity above the background levels existing before the disaster [59]. Therefore, it remains possible that the SV Cycloclasticus observed in our pentane incubations was poised to bloom five years following the spill due to some form of memory effect from the large influx of biomass caused by the disaster.

### Hydrocarbon metabolism across *Cycloclasticus*

To understand how hydrocarbon metabolic capability within Cycloclasticus relates to ecological and evolutionary patterns, we reconstructed *Cycloclasticus* metagenomes from various environments using publicly available datasets (Supplementary Dataset S4). This effort resulted in eight high-quality MAGs with completion of >80% and <2% redundancy. These eight MAGs are in addition to the five pentane MAGs and the one DWH-MAG, and include one from the uncontaminated North Sea “NS_1”, six from a coastal salt marsh in Skidaway Island, Georgia, “CSM_1”, “CSM_2”, “CSM_3”, “CSM_4”, “CSM_5”, and “CSM_6”, and one metagenome from coastal seawater near Pivers Island, North Carolina “CSW_1”. The 14 genomes reconstructed for this study, along with other publicly available genomes, were used to form a phylogenomic tree of all *Cycloclasticus* (Fig. 5a). Each genome was then scanned for hydrocarbon-related pathways of interest and other metabolic functions related to energy generation (Fig. 5b).

From this analysis, we observe distinct metabolic strategies by each major clade within the *Cycloclasticus* genera (Fig. 5b). Notably, all cultivated *Cycloclasticus* are very closely related to each other (Fig. 5a, bottom clade, denoted ‘isolates’). We found no evidence of genes for consuming short-chain alkanes within this clade. This is a major bias in our understanding of *Cycloclasticus*, especially notable because all other *Cycloclasticus* genomes analyzed contained pHMO genes. We also observe two water-column clades from uncontaminated seawater that harbor diverse pathways for short-chain and long-chain alkanes, as well as near-complete pathways for naphthalene and xylene, and complete pathways for toluene and benzene consumption. Altogether, we find a minimum of seven clades within the *Cycloclasticus*, seemingly unified as marine organisms that grow from aqueous soluble hydrocarbons. One key factor distinguishing the clades is the evolved preference to access certain classes of aqueous soluble hydrocarbons and not others.

## Conclusion

Our study provides genomic and proteomic evidence for pentane metabolism by free-living members of the *Cycloclasticus* genus in contrasting oceanic regimes, one with prolific natural seep influence and another far removed from prolific seepage. By comparing *Cycloclasticus* genomes and metagenomes, we show that the hydrocarbon metabolism within this genus is not limited to PAH degradation, with genomic variability enabling different ecotypes to access different ecological niches and structural classes of hydrocarbons. The apparent commonality among *Cycloclasticus* is not the ability to consume aromatic hydrocarbons, as the genus name suggests, but rather a metabolic specialization among the subset of hydrocarbons that exhibit aqueous solubility in marine settings.

Our results further expand on previous findings, illuminating the contrasting strategies between cultivated members of *Cycloclasticus* relying solely on aromatic hydrocarbons and mussel and sponge symbionts primarily consuming short-chain alkanes. We identify distinct clades of free-living *Cycloclasticus* that further expand on these contrasting specializations: a seep variant clade that selectively targets short-chain alkanes using pHMO, and an open ocean variant clade that exhibits broader hydrocarbon versatility and the ability for anaerobic metabolism. A versatile metabolism could benefit *Cycloclasticus* in the ephemeral natural seep environment or in accessing non-seep hydrocarbon sources.

The specialization of SV *Cycloclasticus* on short-chain alkanes is more perplexing because of the apparent ecological risk. The apparent strategy of these SV *Cycloclasticus* requires a consistent supply of short-chain alkanes for growth, which are supplied almost exclusively from petroleum-rich environments such as seeps or spills. Given the geographic constraints on petroleum seepage and the ephemeral nature of discharge, it is perhaps surprising that specialist bacterioplankton have evolved into this niche. Nonetheless, the results of our experiments and the evident success of SV *Cycloclasticus* during DWH indicate that such specialization results in a successful ecological strategy. The success of SV *Cycloclasticus* is likely related to rapid cellular respiration that enables competitive growth upon exposure to the substrate.

The insight gained from this work provides a new vantage for considering the deep ocean microbial response to hydrocarbon discharge during DWH. SV *Cycloclasticus* was well adapted to bloom in response to the massive intrusions of aqueous-soluble *n*-alkanes that accompanied this event. The OOV *Cycloclasticus* may have engaged in direct competition by consuming these same compounds, but they may have also accessed other compounds in parallel or in sequence. This work goes beyond the DWH and provides a predictive capacity for understanding the ocean’s response to future industrial incidents on a variety of scales, such as a rupture of a subsea pipeline or the sinking of a tanker vessel carrying gas condensate, light crude oil, or diluted bitumen; or another well blowout.

## Acknowledgments

We want to thank Dr. Alison Buchan, Dr. Andrew Steen, and Dr. Lauren Quigley for their effort in collecting, sequencing, and making data publicly available from the Groves Creek Marsh in Georgia. We thank the R/V Atlantis captain and crew for their support at sea. This work used the Bridges-2 system for bioinformatic analyses, supported by NSF award number OAC-1928147 at the Pittsburgh Supercomputing Center (PSC). We thank David O’Neal and TJ Olesky for their assistance with bioinformatics optimization and support on the Bridges and Bridges-2 systems. This project was supported by NSF grant numbers OCE-1634478, OCE-1756947, and OCE-2126625.

## References

1. Reddy CM, Arey JS, Seewald JS, Sylva SP, Lemkau KL, Nelson RK, et al. Composition and fate of gas and oil released to the water column during the Deepwater Horizon oil spill. Proceedings of the National Academy of Sciences 2012; 109: 20229–20234.

2. Ryerson TB, Camilli R, Kessler JD, Kujawinski EB, Reddy CM, Valentine DL, et al. Chemical data quantify Deepwater Horizon hydrocarbon flow rate and environmental distribution. Proc Natl Acad Sci U S A 2012; 109: 20246–20253.

3. Gros J, Socolofsky SA, Dissanayake AL, Jun I, Zhao L, Boufadel MC, et al. Petroleum dynamics in the sea and influence of subsea dispersant injection during Deepwater Horizon. Proc Natl Acad Sci U S A 2017; 114: 10065–10070.

4. Sikkema J, de Bont JA, Poolman B. Mechanisms of membrane toxicity of hydrocarbons. Microbiol Rev 1995; 59: 201–222.

5. Van Hamme JD, Singh A, Ward OP. Recent Advances in Petroleum Microbiology. Microbiology and Molecular Biology Reviews 2003; 67: 503–549.

6. Hara A, Syutsubo K, Harayama S. Alcanivorax which prevails in oil-contaminated seawater exhibits broad substrate specificity for alkane degradation. Environ Microbiol 2003; 5: 746– 753.

7. Yakimov MM, Timmis KN, Golyshin PN. Obligate oil-degrading marine bacteria. Curr Opin Biotechnol 2007; 18: 257–66.

8. Valentine DL, Mezić I, Maćešić S, Črnjarić-Žic N, Ivić S, Hogan PJ, et al. Dynamic autoinoculation and the microbial ecology of a deep water hydrocarbon irruption. Proc Natl Acad Sci U S A 2012; 109: 20286–91.

9. Mahmoudi N, Robeson MS, Castro HF, Fortney JL, Techtmann SM, Joyner DC, et al. Microbial community composition and diversity in Caspian Sea sediments. FEMS Microbiol Ecol 2015; 91: 1–11.

10. Techtmann SM, Fortney JL, Ayers KA, Joyner DC, Linley TD, Pfiffner SM, et al. The unique chemistry of Eastern Mediterranean water masses selects for distinct microbial communities by depth. PLoS One 2015; 10: 1–22.

11. Liu J, Techtmann SM, Woo HL, Ning D, Fortney JL, Hazen TC. Rapid Response of Eastern Mediterranean Deep Sea Microbial Communities to Oil. Sci Rep 2017; 7: 5762.

12. McGenity TJ, McKew BA, Lea-Smith DJ. Cryptic microbial hydrocarbon cycling. Nat Microbiol 2021; 6: 419–420.

13. Lea-Smith DJ, Biller SJ, Davey MP, Cotton CAR, Sepulveda BMP, Turchyn A V., et al. Contribution of cyanobacterial alkane production to the ocean hydrocarbon cycle. Proc Natl Acad Sci U S A 2015; 112: 13591–13596.

14. Moulin SLY, Beyly-Adriano A, Cuiné S, Blangy S, Légeret B, Floriani M, et al. Fatty acid photodecarboxylase is an ancient photoenzyme that forms hydrocarbons in the thylakoids of algae. Plant Physiol 2021; 186: 1455–1472.

15. Rubin-Blum M, Antony CP, Borowski C, Sayavedra L, Pape T, Sahling H, et al. Short-chain alkanes fuel mussel and sponge Cycloclasticus symbionts from deep-sea gas and oil seeps. Nat Microbiol 2017; 2: 17093.

16. Saunois M, R. Stavert A, Poulter B, Bousquet P, G. Canadell J, B. Jackson R, et al. The global methane budget 2000-2017. Earth Syst Sci Data 2020; 12: 1561–1623.

17. Coulon F, McKew BA, Osborn AM, McGenity TJ, Timmis KN. Effects of temperature and biostimulation on oil-degrading microbial communities in temperate estuarine waters. Environ Microbiol 2007; 9: 177–186.

18. Love CR, Arrington EC, Gosselin KM, Reddy CM, Van Mooy BAS, Nelson RK, et al. Microbial production and consumption of hydrocarbons in the global ocean. Nat Microbiol 2021; 6: 489–498.

19. Jordan SFA, Gräwe U, Treude T, Lee EM, Schneider von Deimling J, Rehder G, et al. Pelagic methane sink enhanced by benthic methanotrophs ejected from a gas seep. Geophys Res Lett 2021; 63: 183–229.

20. Jordan SFA, Gräwe U, Treude T, Lee EM van der, Deimling JS von, Rehder G, et al. Pelagic Methane Sink Enhanced by Benthic Methanotrophs Ejected From a Gas Seep. Geophys Res Lett 2021; 48: e2021GL094819.

21. Button DK, Robertson BR, Lepp PW, Schmidt TM. A small, dilute-cytoplasm, high-affinity, novel bacterium isolated by extinction culture and having kinetic constants compatible with growth at ambient concentrations of dissolved nutrients in seawater. Appl Environ Microbiol 1998; 64: 4467–4476.

22. Chung WK, King GM. Isolation, Characterization, and Polyaromatic Hydrocarbon Degradation Potential of Aerobic Bacteria from Marine Macrofaunal Burrow Sediments and Description of Lutibacterium anuloederans gen. nov., sp. nov., and Cycloclasticus spirillensus sp. nov. Appl Environ Microbiol 2001; 67: 5585–5592.

23. Fernández-Martínez J, Pujalte MJ, García-Martínez J, Mata M, Garay E, Rodríguez-Valera F. Description of Alcanivorax venustensis sp. nov. and reclassification of Fundibacter jadensis DSM 12178T (Bruns and Berthe-Corti 1999) as Alcanivorax jadensis comb. nov., members of the emended genus Alcanivorax. Int J Syst Evol Microbiol 2003; 53: 331–338.

24. Kozich JJ, Westcott SL, Baxter NT, Highlander SK, Schloss PD. Development of a dual-index sequencing strategy and curation pipeline for analyzing amplicon sequence data on the miseq illumina sequencing platform. Appl Environ Microbiol 2013; 79: 5112–5120.

25. Apprill A, Mcnally S, Parsons R, Weber L. Minor revision to V4 region SSU rRNA 806R gene primer greatly increases detection of SAR11 bacterioplankton. Aquatic Microbial Ecology 2015; 75: 129–137.

26. Parada AE, Needham DM, Fuhrman JA. Every base matters: Assessing small subunit rRNA primers for marine microbiomes with mock communities, time series and global field samples. Environ Microbiol 2016; 18: 1403–1414.

27. Pruesse E, Quast C, Knittel K, Fuchs BM, Ludwig W, Peplies J, et al. SILVA: a comprehensive online resource for quality checked and aligned ribosomal RNA sequence data compatible with ARB. Nucleic Acids Res 2007; 35: 7188–7196.

28. Glöckner FO, Yilmaz P, Quast C, Gerken J, Beccati A, Ciuprina A, et al. 25 years of serving the community with ribosomal RNA gene reference databases and tools. J Biotechnol. 2017. Elsevier., 261: 169–176

29. Bolger AM, Lohse M, Usadel B. Trimmomatic: a flexible trimmer for Illumina sequence data. Bioinformatics 2014; 30: 2114–2120.

30. Joshi N, Fass J. Sickle: A sliding-window, adaptive, quality-based trimming tool for FastQ files (Version 1.33) [Software]. *Available at* https://github.com/najoshi/sickle 2011; 2011.

31. Nurk S, Meleshko D, Korobeynikov A, Pevzner PA. metaSPAdes: a new versatile metagenomic assembler. Genome Res 2017; 27: 824–834.

32. Gurevich A, Saveliev V, Vyahhi N, Tesler G. QUAST: quality assessment tool for genome assemblies. Bioinformatics 2013; 29: 1072–1075.

33. Langmead B, Salzberg SL. Fast gapped-read alignment with Bowtie 2. Nature Methods 2012 9:4 2012; 9: 357–359.

34. Li H, Handsaker B, Wysoker A, Fennell T, Ruan J, Homer N, et al. The Sequence Alignment/Map format and SAMtools. Bioinformatics 2009; 25: 2078–2079.

35. Eren AM, Esen ÖC, Quince C, Vineis JH, Morrison HG, Sogin ML, et al. Anvi’o: an advanced analysis and visualization platform for ‘omics data. PeerJ 2015; 3: e1319.

36. Kim D, Song L, Breitwieser FP, Salzberg SL. Centrifuge: rapid and sensitive classification of metagenomic sequences. Genome Res 2016; 26: 1721–1729.

37. Parks DH, Imelfort M, Skennerton CT, Hugenholtz P, Tyson GW. CheckM: assessing the quality of microbial genomes recovered from isolates, single cells, and metagenomes. Genome Res 2015; 25: 1043–1055.

38. Chaumeil P-A, Mussig AJ, Hugenholtz P, Parks DH. GTDB-Tk: a toolkit to classify genomes with the Genome Taxonomy Database. Bioinformatics 2020; 36: 1925–1927.

39. Rodriguez-R LM, Konstantinidis KT. The enveomics collection: a toolbox for specialized analyses of microbial genomes and metagenomes. PeerJ Prepr 2016; 4: e1900v1.

40. Kang DD, Li F, Kirton E, Thomas A, Egan R, An H, et al. MetaBAT 2: An adaptive binning algorithm for robust and efficient genome reconstruction from metagenome assemblies. PeerJ 2019; 2019: e7359.

41. Edgar RC. MUSCLE: a multiple sequence alignment method with reduced time and space complexity. BMC Bioinformatics 2004 5:1 2004; 5: 1–19.

42. Capella-Gutiérrez S, Silla-Martínez JM, Gabaldón T. trimAl: a tool for automated alignment trimming in large-scale phylogenetic analyses. Bioinformatics 2009; 25: 1972–1973.

43. Stamatakis A. RAxML version 8: a tool for phylogenetic analysis and post-analysis of large phylogenies. Bioinformatics 2014; 30: 1312–1313.

44. FigTree. http://tree.bio.ed.ac.uk/software/figtree/. Accessed 14 Oct 2021.

45. Timmins-Schiffman E, May DH, Mikan M, Riffle M, Frazar C, Harvey HR, et al. Critical decisions in metaproteomics: achieving high confidence protein annotations in a sea of unknowns. The ISME Journal 2017 11:2 2016; 11: 309–314.

46. Eng JK, Jahan TA, Hoopmann MR. Comet: An open-source MS/MS sequence database search tool. Proteomics 2013; 13: 22–24.

47. Nesvizhskii AI, Keller A, Kolker E, Aebersold R. A statistical model for identifying proteins by tandem mass spectrometry. Anal Chem 2003; 75: 4646–4658.

48. Ma K, Vitek O, Nesvizhskii AI. A statistical model-building perspective to identification of MS/MS spectra with PeptideProphet. BMC Bioinformatics 2012; 13 **Suppl 1**: 1–17.

49. Aroney, Samuel T N; Newell, Rhys; Nissen, J; Camargo, Antonio; Tyson, Gene; Woodcroft B. CoverM: Read coverage calculator for metagenomics. 2024.

50. Jordan SFA, Treude T, Leifer I, Janßen R, Werner J, Schulz-Vogt H, et al. Bubble-mediated transport of benthic microorganisms into the water column: Identification of methanotrophs and implication of seepage intensity on transport efficiency. Sci Rep 2020; 10: 1–15.

51. Vogel AL, Thompson KJ, Straub D, Musat F, Gutierrez T, Kleindienst S. Genetic redundancy in the naphthalene-degradation pathway of Cycloclasticus pugetii strain PS-1 enables response to varying substrate concentrations. FEMS Microbiol Ecol 2024; 100: 60.

52. White H, Huber C, Feicht R, Simon H. On a reversible molybdenum-containing aldehyde oxidoreductase from Clostridium formicoaceticum. Arch Microbiol 1993; 159: 244–249.

53. Wang W, Wang L, Shao Z. Polycyclic aromatic hydrocarbon (PAH) degradation pathways of the obligate marine PAH degrader Cycloclasticus sp. strain P1. Appl Environ Microbiol 2018; 84.

54. Moran MA, Buchan A, González JM, Heidelberg JF, Whitman WB, Kiene RP, et al. Genome sequence of Silicibacter pomeroyi reveals adaptations to the marine environment. Nature 2005 432:7019 2004; 432: 910–913.

55. Valentine DL, Kessler JD, Redmond MC, Mendes SD, Heintz MB, Farwell C, et al. Propane respiration jump-starts microbial response to a deep oil spill. Science 2010; 330: 208–11.

56. Redmond MC, Valentine DL. Natural gas and temperature structured a microbial community response to the Deepwater Horizon oil spill. Proc Natl Acad Sci U S A 2012; 109: 20292–20297.

57. Kessler JD, Valentine DL, Redmond MC, Du M, Chan EW, Mendes SD, et al. A Persistent Oxygen Anomaly Reveals the Fate of Spilled Methane in the Deep Gulf of Mexico. Science (1979) 2011; 331: 312–315.

58. Rivers AR, Sharma S, Tringe SG, Martin J, Joye SB, Moran MA. Transcriptional response of bathypelagic marine bacterioplankton to the Deepwater Horizon oil spill. The ISME Journal 2013 7:12 2013; 7: 2315–2329.

59. Rogener MK, Bracco A, Hunter KS, Saxton MA, Joye SB. Long-term impact of the Deepwater Horizon oil well blowout on methane oxidation dynamics in the northern Gulf of Mexico. Elem Sci Anth 2018; 6: 73.

60. MacDonald IR, Garcia-Pineda O, Beet A, Daneshgar Asl S, Feng L, Graettinger G, et al. Natural and unnatural oil slicks in the Gulf of Mexico. J Geophys Res Oceans 2015; 120: 8364–8380.

