## Supplementary Information for "Hydrocarbon Metabolism and Petroleum Seepage as Ecological and Evolutionary Drivers for *Cycloclasticus*"

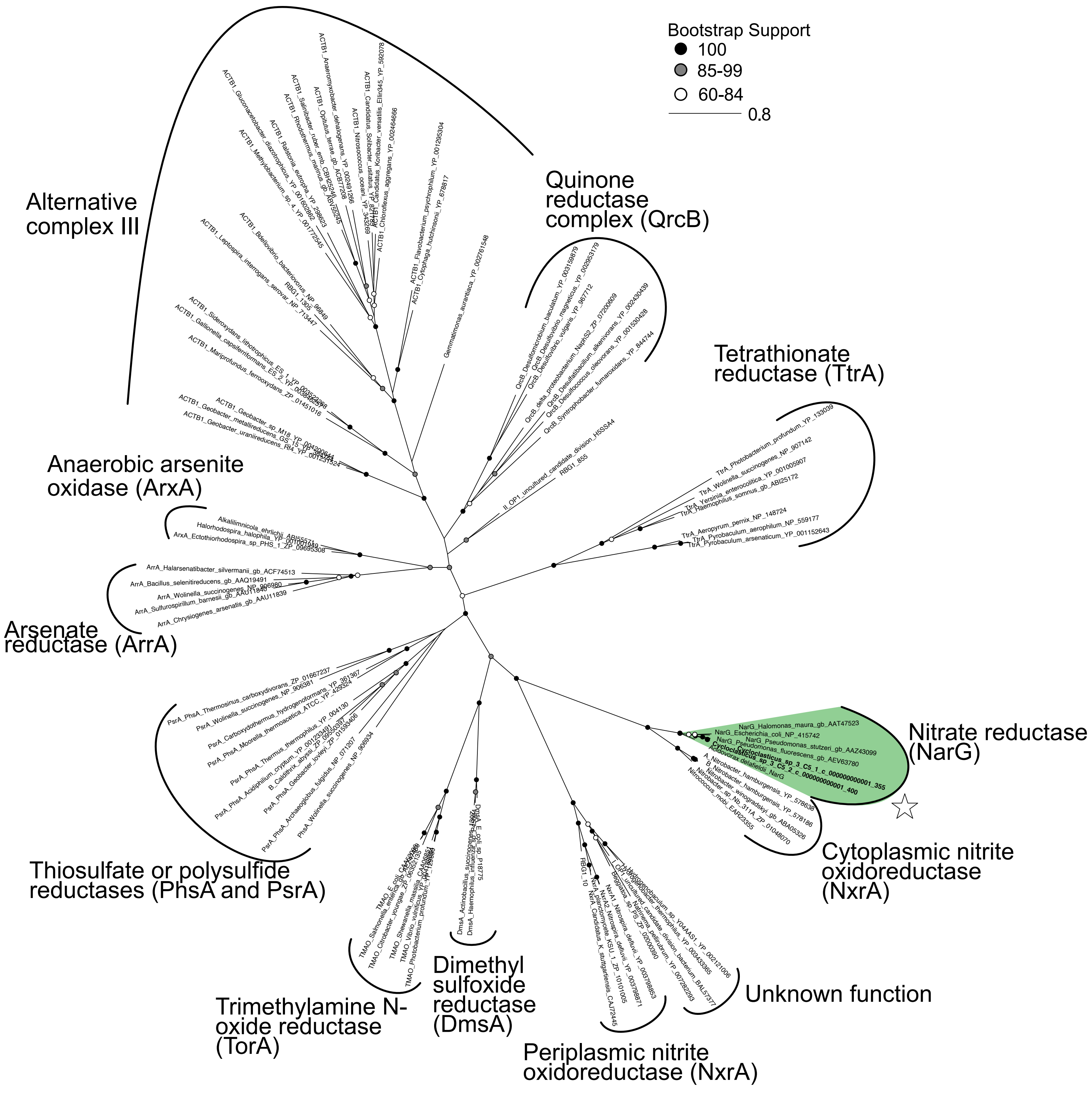


**Supplemental Data Figure S1.** Maximum-likelihood phylogenetic tree with scale bar of substitutions per site of DMSO reductase superfamily modeled after [1]. NarG sequences from Cycloclasticus OOV variant in bold and noted with star symbol.

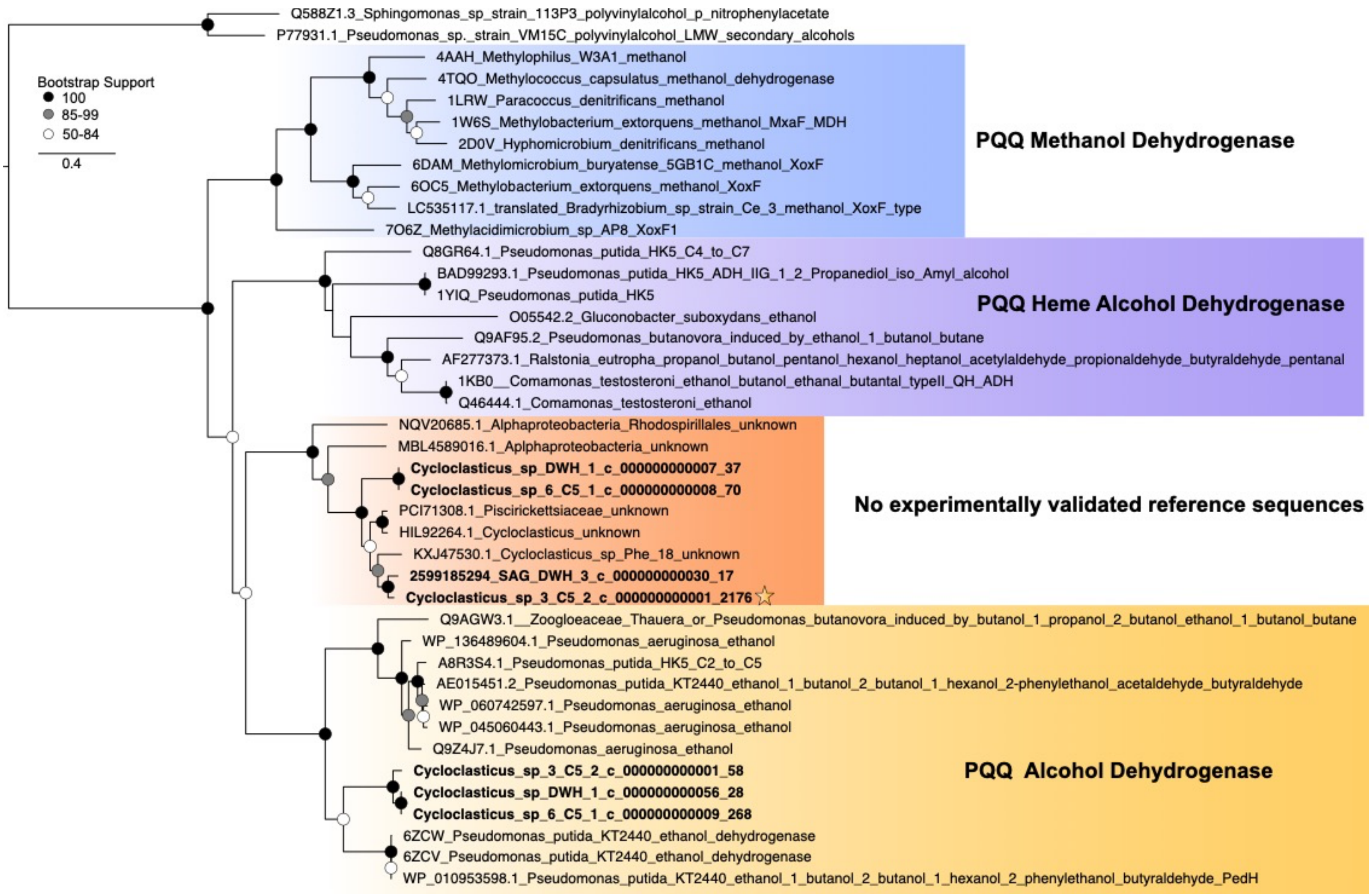


**Supplemental Figure S2.** Maximum-likelihood phylogenetic tree of PQQ-dependent alcohol dehydrogenase sequences rooted to polyvinylalcohol reference sequences. Experimentally validated reference sequences are annotated with known substrate activity. Sequence denoted with a star was observed in proteomic samples from pentane enrichment. No experimentally validated reference sequences present in the orange clade.

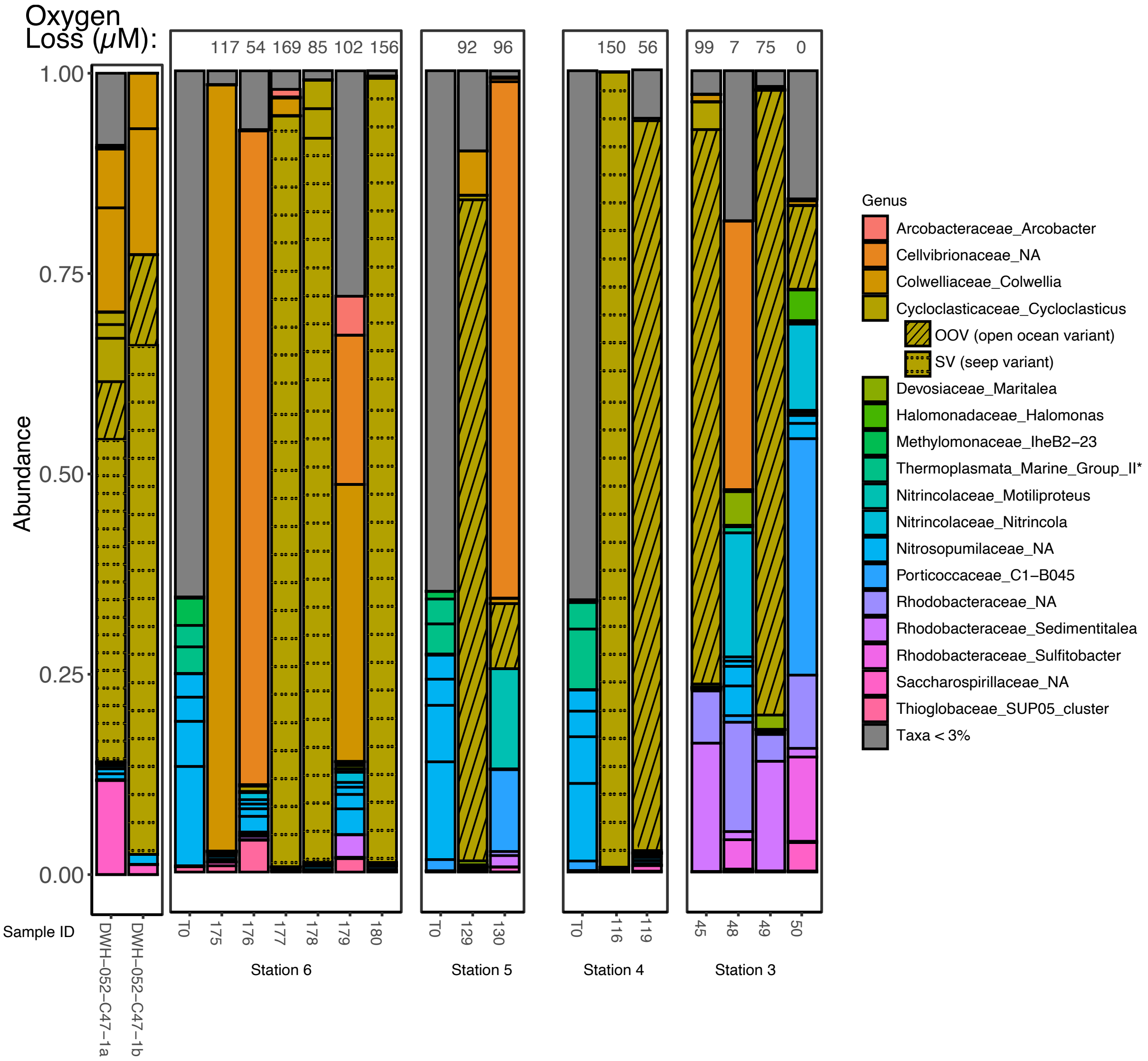


**Supplemental Figure S3.** 16S rRNA community analysis of V4 region. Sample 119, 48, and 50 are “non-bloom” samples.

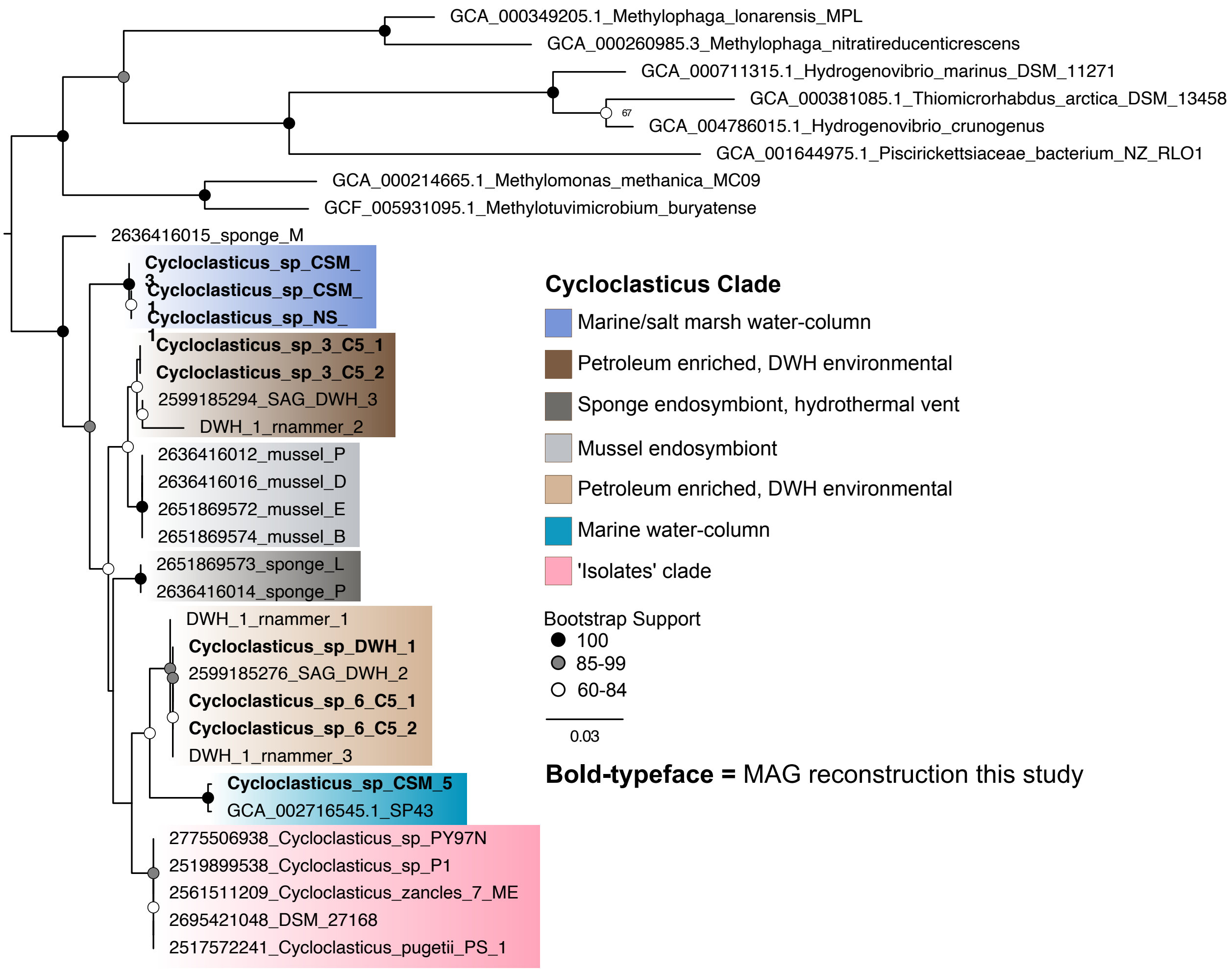


**Supplemental Figure S4.** Maximum-likelihood phylogenetic tree of 16S rRNA rooted to representatives from *Piscirickettsiaceae* and *Methylococcaceae* family. "_rnammer" denotes 16S sequences recovered from assembled contigs prior to binning for the DWH sample. Alignment includes full length 16S rRNA gene 1,547 amino acid residues.

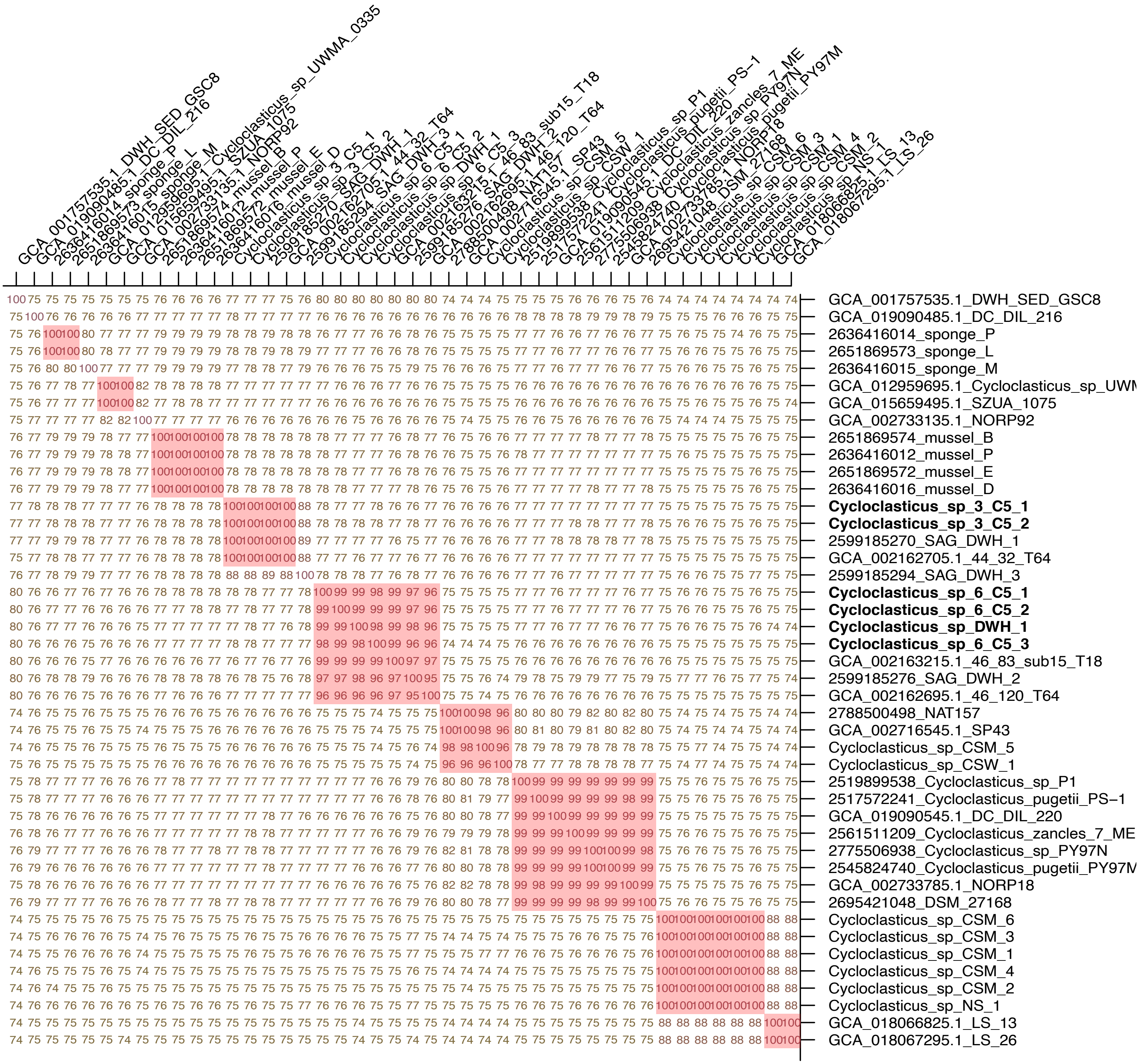


**Supplemental Figure S5.** Average nucleotide identity of *Cycloclasticus* genomes. SV and OOV are highlighted in bold.

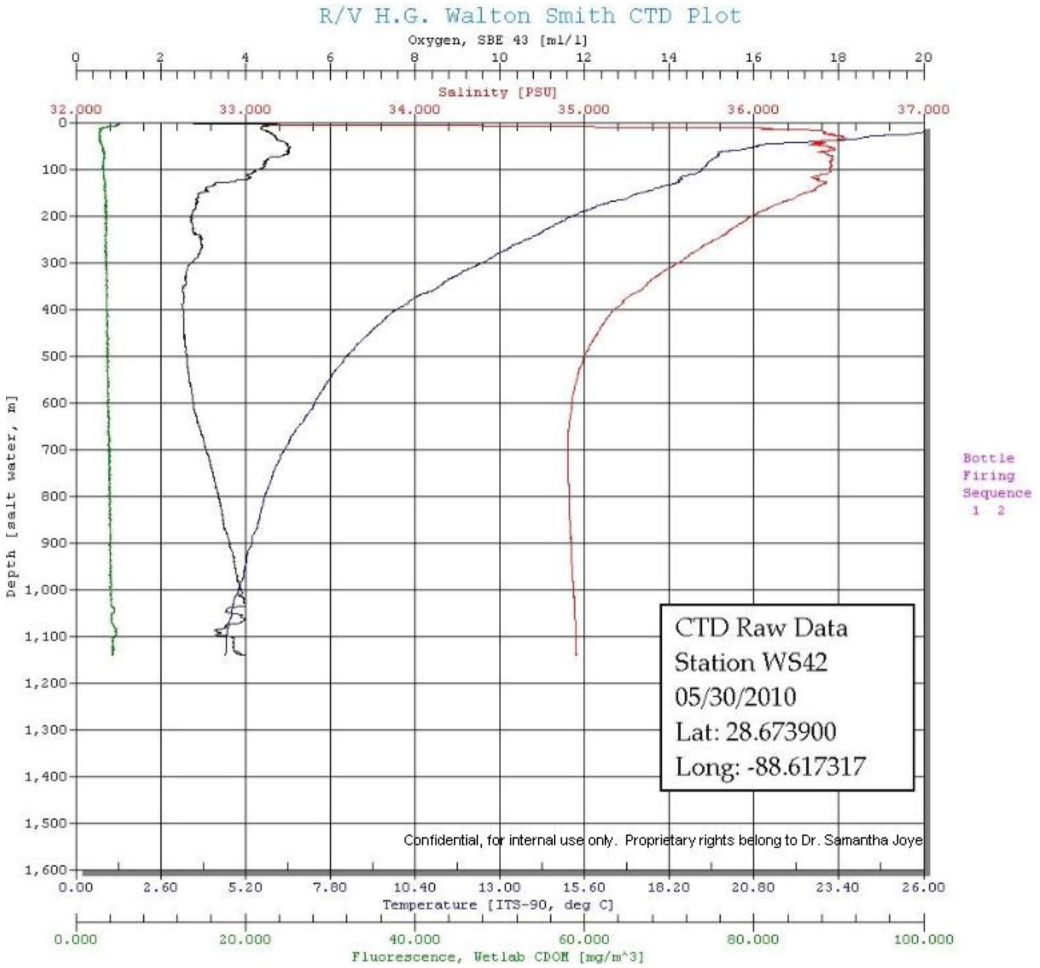


**Supplemental Figure S6.** CTD profile from cast during the DWH event. DWH sample analyzed in this study was collected at 1,090m by Dr. Molly Redmond. [2]

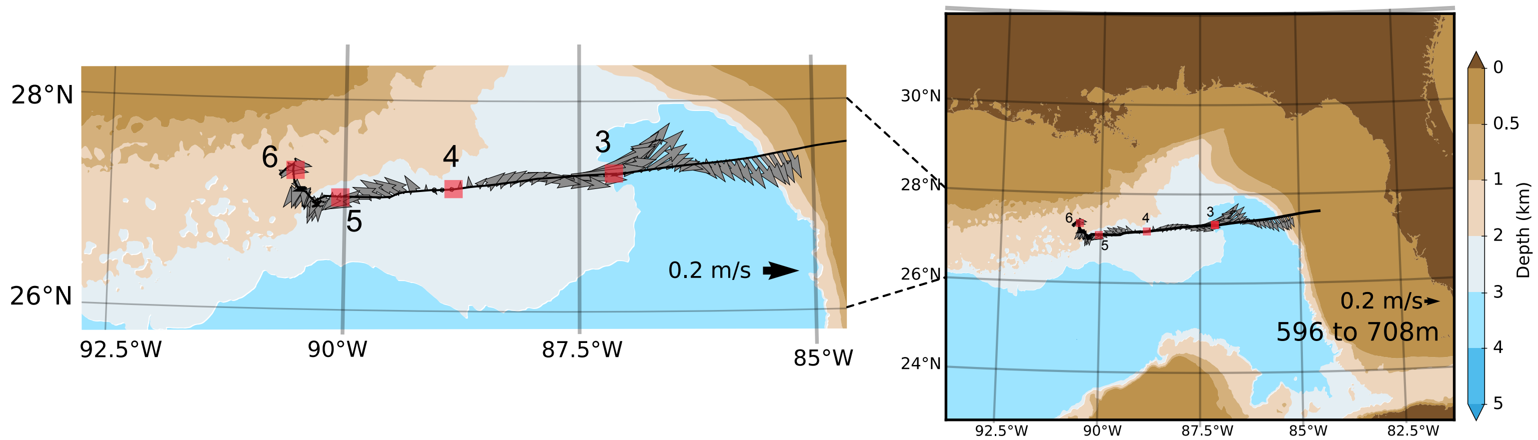


**Supplemental Data Figure S7.** ADCP data showing deep ocean currents over the course of the GOM sampling expedition (RV Atlantis, 2015).

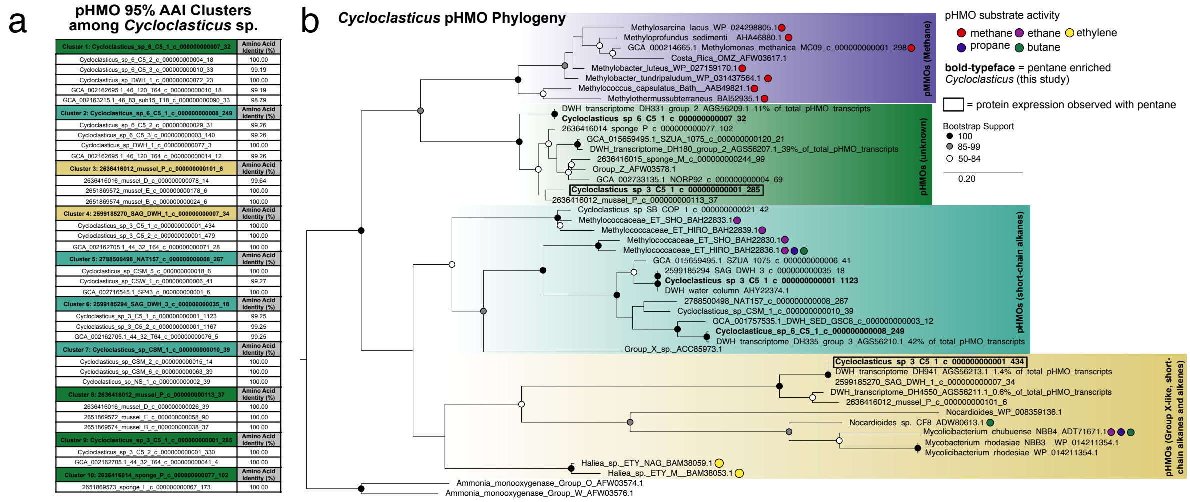


**Supplemental Data Figure S8.** Phylogeny of pHMO subunit A protein sequences. **a** Description of pHMO clusters containing 95% amino acid identity. Amino acid identity to the representative sequence is noted. The representative sequence for each cluster is colored according to the phylogenetic placement. **b** Maximum likelihood tree of representative sequences is drawn to scale, with branch lengths representing the number of substitutions per site. Bootstrap values below 50% are not shown. Each major clade is color-coded for readability with purple representing pMMOs with activity on methane, the green clade is "unknown" and lacks any known substrate specificity, the cyan clade represents group X pHMOs (ethane/ethylene, propane, and butane activity), and the yellow clade are group-X like (ethane, propane, and butane activity). Sequences from the pentane-enriched *Cycloclasticus* MAGs are in bold, boxed values indicate pHMOs detected in proteomic data.

References

1. Castelle CJ, Hug LA, Wrighton KC, Thomas BC, Williams KH, Wu D, et al. Extraordinary phylogenetic diversity and metabolic versatility in aquifer sediment. *Nat Commun* 2013; **4**.
2. National Oceanic and Atmospheric Administration *R/V Walton Smith CTD profile data*. Available at https://www.ncei.noaa.gov/data/oceans/DeepwaterHorizon/Ship/Walton_Smith/ORR/Cruise_01/CTD/Products/. Accessed 10 Jun 2024.
